# Natural chromatin is heterogeneous and self associates *in vitro*

**DOI:** 10.1101/139543

**Authors:** Shujun Cai, Yajiao Song, Chen Chen, Jian Shi, Lu Gan

## Abstract

The 30-nm fiber is commonly found in oligonucleosome arrays *in vitro* but rarely found in chromatin within nuclei. To determine how chromatin high-order structure is controlled, we used cryo-ET to study the undigested natural chromatin released from cells that do not have evidence of 30-nm fibers *in vivo*: picoplankton and yeast. In the presence of divalent cations, most of the chromatin from both organisms is compacted into a large mass. Rare irregular 30-nm fibers do form at the periphery of this mass, some of which include face-to-face interactions. In the absence of divalent cations, picoplankton chromatin decondenses into open zigzags. By contrast, yeast chromatin mostly remains compact with looser nucleosome packing, even after treatment with histone-deacetylase inhibitor. The 3-D configuration of natural chromatin is therefore sensitive to the local environment, but generally nonpermissive of regular motifs, even at the level of oligonucleosomes.

## INTRODUCTION

In eukaryotic cells, chromatin is important in nuclear processes like transcription and replication. Chromatin structure is dictated by the 3-D relationship between nucleosomes (∼ 146 bp of DNA wrapped around 8 histones). Chromatin can adopt open or compact higher-order structures, influenced by factors like histone post-translational modifications and interactions with chromatin “architectural” proteins (Jenuwein and Allis, 2001; McBryant et al., 2006). Chromatin higher-order structure in turn controls the accessibility to the DNA for both transcription and replication (Collins et al., 2002; Tse et al., 1998).

Chromatin structure has been studied extensively at nucleosome level. Nucleosomes connected by linker DNA are described as “beads-on-a-string” (Olins and Olins, 1974). The beads-on-a-string is hypothesized to be compacted into a helical nucleosome arrangement called the “30-nm fiber”, which is proposed to be either a one-start helix or two-start zigzag (Finch and Klug, 1976; Woodcock et al., 1984). However, more than one form of nucleosome packing might also coexist in the same 30-nm fiber (Grigoryev et al., 2009).

The diversity of chromatin structures has lead to inconsistent terminology. For the sake of clarity, we define a chromatin “mass” as a compact body with irregularly packed chromatin; a “regular 30-nm fiber” as an ordered compact structure approximately 30-nm wide; an “irregular 30-nm fiber” as a fiber-like structure with few repeated nucleosome motifs and of variable width; an “open zigzag” as a more extended structure with resolvable linker DNA and few nucleosome-nucleosome interactions; and the beads-on-a-string as a chain of nucleosomes in which the angle between the entering and exiting linker DNA is nearly 180° (Fig. 1A).

**Figure 1.**
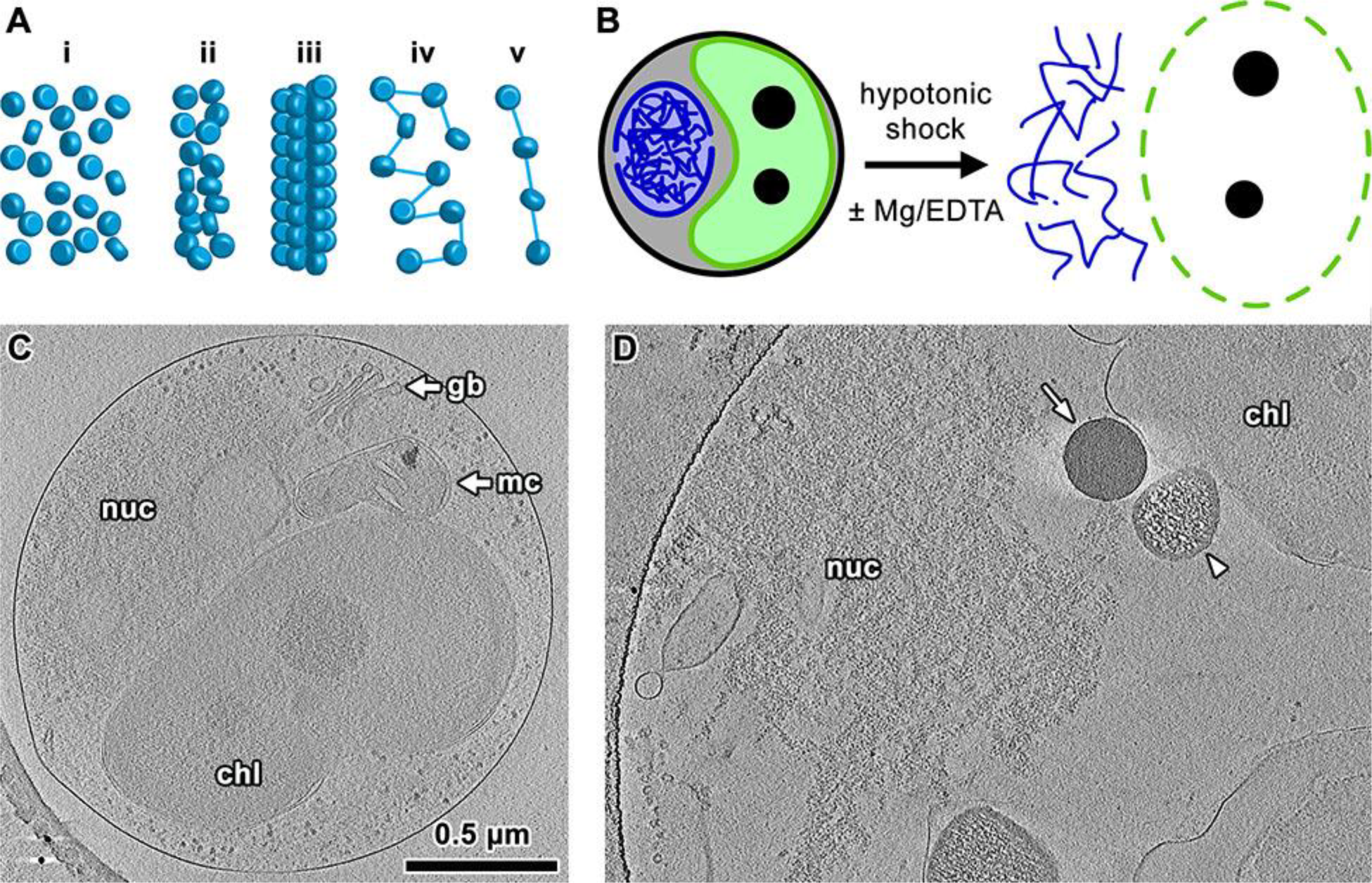
Strategy for 3-D analysis of natural chromatin. (A) Possible nucleosome (blue discs) arrangements, including (i) chromatin masses, (ii) irregular 30-nm fibers, (iii) regular 30-nm fibers, (iv) open zigzags, (v) beads on a string. Blue lines: linker DNA, depicted based on experimental observations. (B) The plasma membrane is ruptured by hypotonic shock, exposing the cellular contents to the final ionic conditions (with or without Mg^2+^ or EDTA added). The chloroplast (green) and nucleus (blue structure) are the two largest organelles in picoplankton. The dashed lines indicate the disruption of the plasma (black), chloroplast (green), and nuclear (blue) membranes. (C) Tomographic slice (30 nm) showing an intact early-interphase picoplankton cell. The largest organelles are labelled: chloroplast (chl); nucleus (nuc); Golgi body (gb); mitochondrion (mc). The lower left corner shows the curved edge of the carbon support and two gold fiducials (dark puncta). Note that as previously observed, the nucleus is frequently found without an intact nuclear envelope even in interphase. (D) Tomographic slice (30 nm) of a lysed picoplankton cell in 1 mM Mg^2+^, at the same scale as panel C. The arrow points to the radiation-tolerant granule from the chloroplast and the arrowhead shows the chloroplast-derived granule that is sensitive to electron beam damage. The remnants of the nucleus (nuc) and chloroplast (chl) are labelled. The lysed cell is flattened by surface tension (after blotting), causing all components components to spread out. This spreading allows the ice to be much thinner resulting in higher image contrast than an intact cell.

Most of our knowledge of chromatin structure comes from studies of dilute solutions of short nucleosomes chains. These studies have systematically tested the effects of fixation with glutaraldehyde (Athey et al., 1990), recombinant histones (Luger et al., 1997b), linker histones (Routh et al., 2008), artificially selected “Widom 601” sequences that have a high affinity for histones (Dorigo et al., 2004), and ionic conditions (Dorigo et al., 2003; Huynh et al., 2005). Under special conditions, such chromatin constructs can form highly regular 30-nm fibers that can be studied at high resolution (Song et al., 2014).

Chromatin structure has also been studied *in situ* in a life-like frozen-hydrated state, either in 2-D by cryo-EM projection imaging or in 3-D by electron cryotomography (cryo-ET). These studies are much rarer because it is challenging to thin and then image the cells at cryogenic temperatures. Three groups found no evidence of regular 30-nm fibers in mammalian cells (McDowall et al., 1986; Eltsov et al., 2008; Eltsov et al., 2014). We found that in the single-celled organisms *Ostreococcus tauri* (a eukaryotic picoplankton; herein called picoplankton) and *Saccharomyces cerevisiae* (a budding yeast; herein called yeast), there was also no evidence of regular 30-nm fibers (Gan et al., 2013; Chen et al., 2016). In contrast, cryo-EM studies have detected regular 30-nm fibers in the isolated nuclei of both starfish sperm (Woodcock, 1994) and chicken erythrocytes (Scheffer et al., 2011). These and other studies challenge whether 30-nm fibers is the best model of chromatin within somatic cells (Hansen, 2012; Nishino et al., 2012; van Holde and Zlatanova, 1995).

The different conclusions between *in vitro* and *in vivo* chromatin studies raise the question: can the chromatin of cells that do not show evidence of 30-nm fibers *in vivo* be made to adopt a 30-nm-fiber structure *in vitro*? To address this question, we used cryo-ET to study the 3-D organization of undigested natural chromatin from picoplankton and yeast. In the presence of even traces of divalent cations, most of the chromatin in both organisms compacts into large masses. Some of the chromatin does form 30-nm fibers, predominantly of the irregular variety. In the absence of divalent cations, picoplankton chromatin decondenses into open zigzags, but yeast chromatin remains either in a mass or folded as irregular 30-nm fibers. To better understand how these nucleosomes interact, we classified the chromatin in both 2-D and 3-D, and found that there is no dominant higher-order packing motif, but the orientation of the DNA at the core-particle entry/exit points is not too variable. Therefore, the higher-order structure of natural chromatin is sensitive to environmental conditions, but this sensitivity varies by species.

## RESULTS

### Release of natural picoplankton chromatin

To study the higher-order structure of natural picoplankton chromatin *in vitro*, we lysed cells in hypotonic buffer on ice either with or without divalent cations (Fig. 1B). This treatment released all cellular contents, including the chromatin, and is expected to produce thinner plunge-frozen samples that generate higher-contrast cryotomograms. Compared to intact cells (Fig. 1C), the lysed cells’ contents spread over a much larger area and indeed allowed the ice to be thinner, resulting in higher-contrast tomograms (Fig. 1D). The majority of the densities came from remnants of the two largest organelles – the nucleus and the chloroplast. To keep the chromatin as intact as possible, we did not perform any isolation procedures or nuclease digestion.

### Picoplankton chromatin forms masses and irregular 30-nm fibers only in the presence of divalent cations

Previous studies have shown that short nucleosome arrays are sensitive to divalent cation concentration (Bednar et al., 1998). To test if divalent cations can induce 30-nm-fibers in undigested picoplankton chromatin, we lysed and imaged picoplankton in the presence of 1 mM Mg^2+^. Most of the chromatin in these conditions was compacted into a large thick mass, but a small portion of the chromatin did form irregular 30-nm fibers (Figs. 2A and S1). The contrast was too low, however, to reveal more details. To determine how much the compaction and 30-nm fibers depended on Mg^2+^, we lysed cells without additional divalent cations (Fig. 2B). Under this condition, most of the chromatin still remained associated as a large compact mass, but with more short irregular 30-nm fibers visible (Figs. 2E,F and S2). As a consequence of the ice thickness and dense nucleosome packing, the contrast was still too low to permit the unambiguous visualization of individual nucleosomes in most tomograms. In rare examples, we found some small clusters of nucleosomes packed in regular face-to-face interactions (Figs. S2G-J).

**Figure 2.**
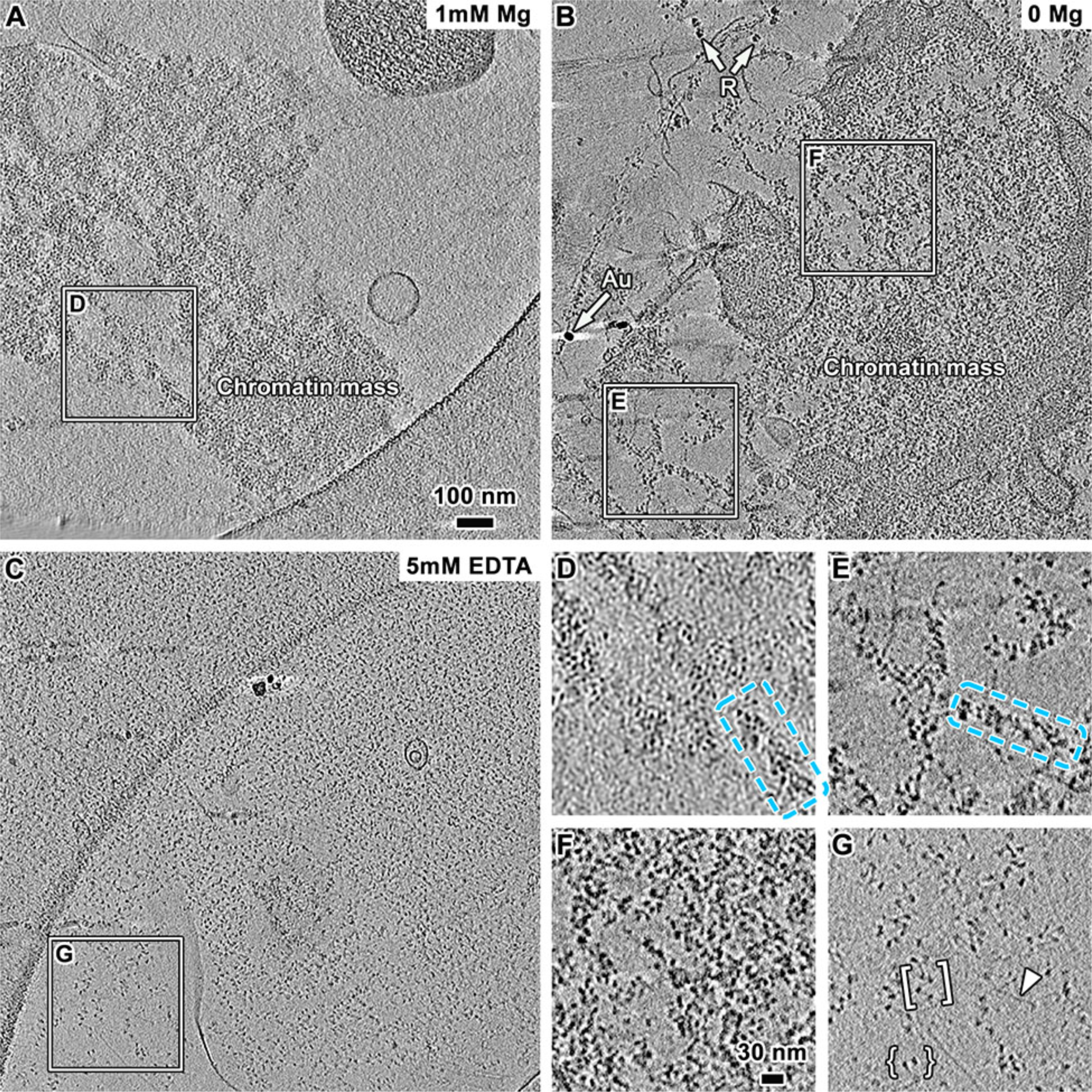
Picoplankton chromatin can form masses and irregular 30-nm fibers in the presence of divalent cations. Tomographic slices (30 nm) of picoplankton cell lysates in buffer supplemented with (A) 1 mM Mg^2+^, (B) no additional divalent cations, and (C) 5 mM EDTA. The dense round structures are gold fiducials (Au in panel B); the gently curved structures in A and C are from the edges of the carbon support film. A couple of ribosomes (R) are indicated with arrows in (B). Positions boxed in A - C are enlarged 2-fold, showing irregular 30-nm fibers (D-F), open zigzags (G, bracket), and a couple of nucleosomes (G, curly brackets). A long stretch of naked DNA is indicated with an arrowhead in panel G. More examples of chromatin subjected to each condition are shown in Figs S2 - S4.

To determine if picoplankton chromatin masses and irregular 30-nm fibers depend on divalent cations, we chelated the residual divalent cations by lysing cells in the presence of 5 mM Ethylenediaminetetraacetic Acid (EDTA) (Fig. 2C, S3). The EDTA treatment was effective because ribosomes, which require magnesium for stability, were absent (Chao, 1957). In picoplankton samples treated this way, we did not find any evidence of 30-nm fibers or chromatin masses. Instead, all of the chromatin was decondensed into an open zigzag-like motif with individual nucleosomes and naked DNA visible (Fig. 2G). This large-scale change of chromatin higher-order structure demonstrates that picoplankton chromatin higher-order structure is very sensitive to divalent cation concentration.

### Yeast chromatin packing is less sensitive to divalent cations than picoplankton

Picoplankton and yeast chromatin do not show evidence of 30-nm fibers *in vivo* (Gan et al., 2013; Chen et al., 2016). To test whether the chromatin higher-order structure of these two organisms is conserved *in vitro*, we imaged yeast chromatin in cell lysate. Yeast chromatin was more difficult to locate in these samples because there was no chromatin-proximal high-contrast structure like the chloroplast. To overcome this problem, we first isolated and then lysed the nuclei in hypotonic buffer without additional divalent cations (Fig. S4). This strategy made it straightforward to locate the chromatin because it was now the most abundant material on the grid.

Like in picoplankton, the majority of yeast chromatin was compacted in a thick mass (Figs. 3A). This chromatin mass had low contrast, which prevented us from visualizing individual nucleosomes. In rare cases, we saw 30-nm fibers at the periphery of the chromatin mass (Fig. 3A). In total, we observed only seven irregular 30-nm fibers in eleven tomograms. These observations show that in the presence of trace amounts of free magnesium, yeast chromatin compacts *in vitro* and can occasionally form irregular 30-nm fibers.

**Fig. 3.**
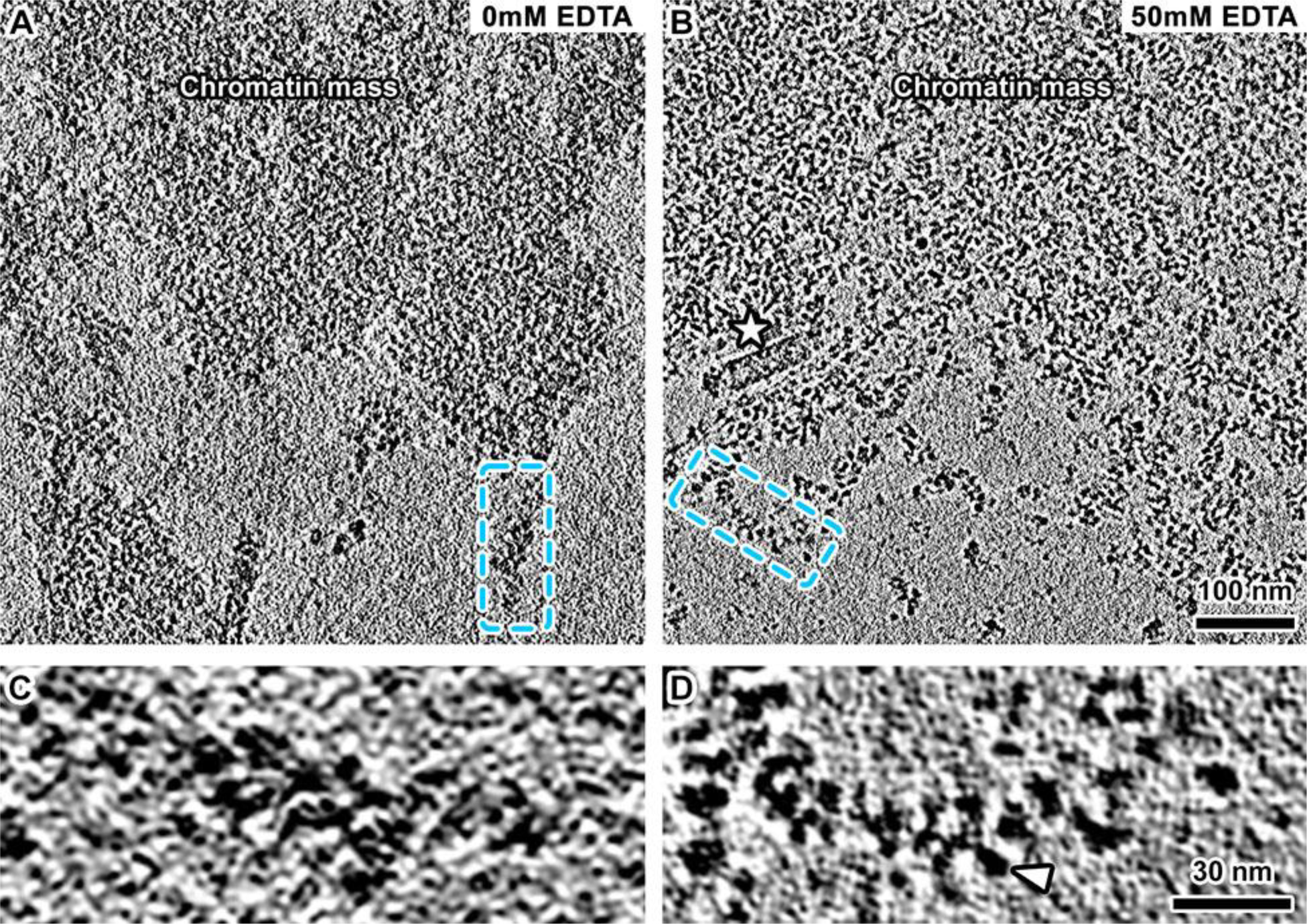
Yeast chromatin forms masses and irregular 30-nm fibers even in the absence of divalent cations. (A) Tomographic slices (30 nm) of yeast nuclei lysate without EDTA. (B) Tomographic slices (30 nm) of yeast nuclei lysate with 50 mM EDTA. Star: microtubule. (C) Enlargement (4-fold) of 30-nm fiber boxed in panel A. (D) Enlargement (4-fold) of an irregular 30-nm fiber in 50 mM EDTA. Arrowhead: nucleosome.

To test if the yeast chromatin organization depends on residual divalent cations, we lysed nuclei in the presence of EDTA. In the 5 mM EDTA-treated sample, we were unable to distinguish most nucleosomes because the chromatin was not dispersed enough (Fig. S5A,C,D). Intact ribosomes could still be found, indicating that trace amounts of divalent cations were still present (Fig. S5A and C). We therefore increased the lysis-buffer EDTA concentration to 50 mM. This new condition was more effective because we could not find any 80S ribosomes (Fig. S5B and E). In the presence of 50 mM EDTA, the chromatin mass became more dispersed (Fig. S6), and unlike in the conditions with less EDTA, chromatin fibers thicker than 50 nm were no longer detectable (Fig. S6). Unlike in picoplankton, in the absence of divalent cations, yeast nucleosomes remained closely packed and were not found to adopt the open zigzags with visible linker DNA (Fig. 3B, more examples in Fig. S5E and F). In summary, natural yeast chromatin can compact into a large mass and form irregular 30-nm fibers even in the absence of divalent cations.

### Histone deacetylation facilitates yeast chromatin compaction

In an early study (Lowary and Widom, 1989), yeast chromatin fragments in low salt formed extended “10-nm filaments”. We did not observe such structures. While the conditions in that study were different from ours (traditional EM vs. cryo-EM; digested vs. undigested chromatin), we sought another explanation for the differences observed. It is known that yeast chromatin is susceptible to deacetylation when cells are disrupted (Waterborg, 2000). To test whether the histone acetylation levels in yeast chromatin *in vitro* were too low to support a more disperse structure like the open zigzag, we treated the intact cells with Trichostatin A (TSA), a histone-deacetylase inhibitor (Bernstein et al., 2000). We then lysed the cells in hypotonic buffer containing both EDTA and TSA. We found that chromatin isolated this way was indeed more disperse (Figs. 4A and B, more examples in Fig. S7). In rare cases, we found open-zigzags (Figs. 4C and D), but we still did not find any structures that resembled a 10-nm filament. These results are nevertheless consistent with a recent study that showed that histone acetylation makes chromatin more deformable (Shimamoto et al., 2017).

**Figure 4.**
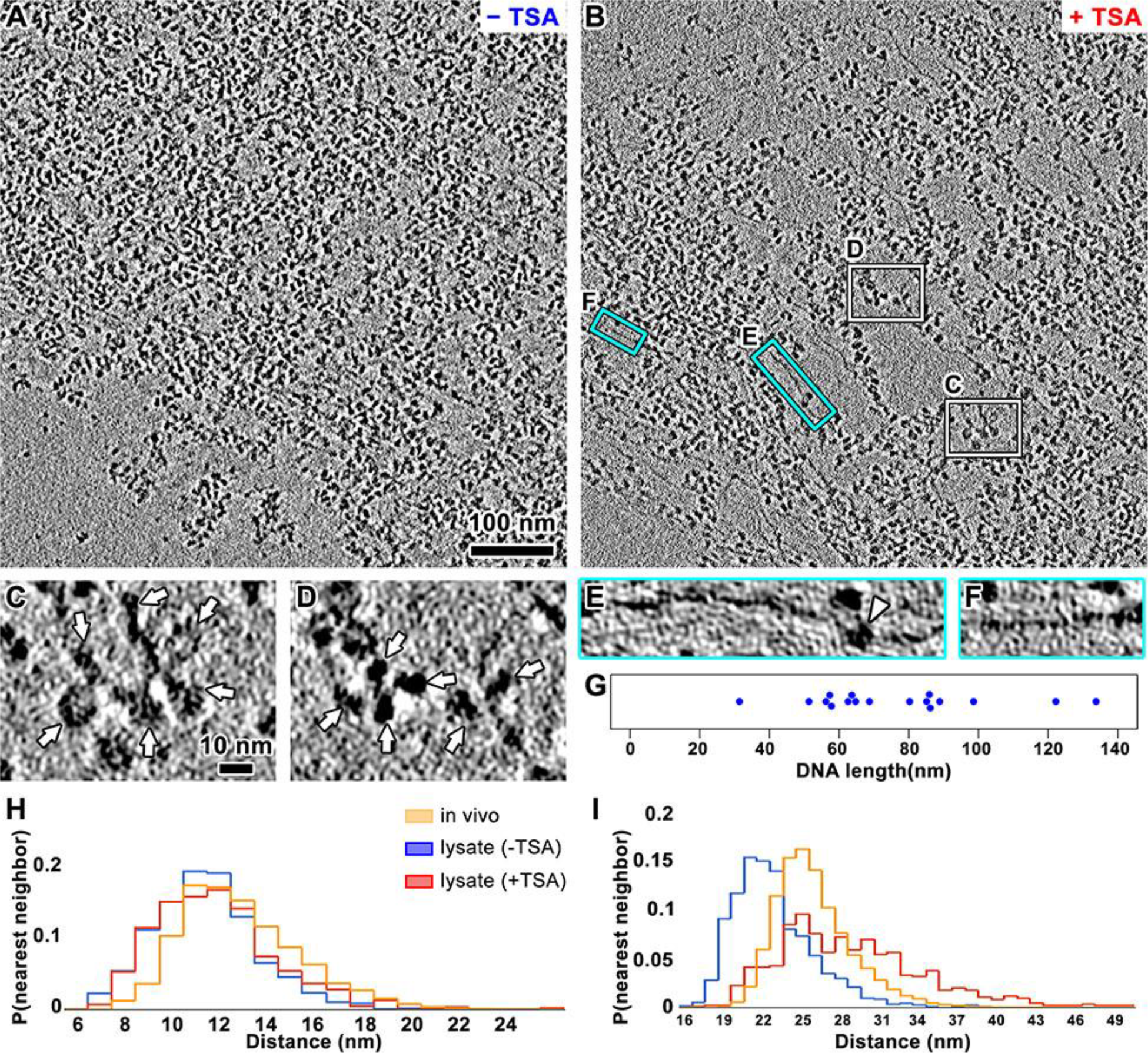
Inhibition of histone deacetylase partially disperses yeast chromatin. (A) Tomographic slice (10 nm) of yeast nuclei lysed in buffer with 50 mM EDTA. (B) Tomographic slice (10 nm) of yeast nuclei lysed in buffer with 50 mM EDTA and 82 µM TSA. (C and D) Four-fold enlargements of positions boxed in white in panel B, showing open zigzags. Arrows: nucleosomes. (E and F) Four-fold enlargements of positions boxed in cyan in panel B, rotated counter clockwise. The naked DNA is 110 nm long in panel E and 55 nm long in panel F. Arrowhead: a non-nucleosome density. (G) tion of naked DNA length in tomograms of (B). (H and I) Histograms of nearest­th nearest-neighbor distances, respectively. The *in vivo* measurements were ing a cryotomogram of sectioned yeast we previously reported (Chen et al., 2016).

To analyze the nucleosome packing more quantitatively, we performed template matching and then analysed both the nearest-neighboring distance (NND) and 10^th^ NND distributions. The NND reports on short-range compaction whereas the 10^th^ NND reports on long-range compaction, e.g., inter-fiber interactions (Fig. S8). As a reference, we also performed this analysis for nucleosome-like densities *in vivo* (Chen et al., 2016). The NND distribution plots of chromatin *in vivo* and both with and without TSA treatment were almost indistinguishable (Fig. 4H). Note, however, that template matching *in vivo* produces more false positives due to lower contrast and the presence of more non-nucleosomal macromolecular complexes. These extra false positives make the *in vivo* NND values lower than they really are. Therefore, nucleosomes are very likely to be packed more loosely *in vivo*. In contrast, the 10^th^ NND distribution of TSA-treated chromatin was longer than chromatin *in vivo*, which was itself longer than untreated chromatin *in vitro* (Fig. 4I). Our NND analysis is consistent with chromatin’s appearance: histone acetylation does not severely perturb short-range interactions, but does affect long-range interactions between groups of nucleosomes.

In the chromatin of TSA-treated yeast, we observed a few stretches of naked DNA between nucleosomes (Fig. 4B,E,F). Nucleosome-free regions are a form of non-nucleosome-associated DNA that are usually longer than one nucleosome’s worth of DNA (Lee et al., 2007; Mavrich et al., 2008; Shivaswamy et al., 2008; Yuan et al., 2005). To test if the long stretches of naked DNA could be nucleosome-free regions, we measured their lengths. We found that most of the naked DNA is between 50 nm and 100 nm long, which is compatible with the absence of one or two nucleosome core particles (each 50 nm long) (Fig. 4G). Our data would also be consistent with the notion that this naked DNA was wrapped around a ‘fragile’ nucleosome or pre-nucleosome *in vivo*, but then lost their histones during nuclear lysis (Fei et al., 2015; Kubik et al., 2015). On some of the naked DNA strands, we found densities that are smaller than a nucleosome (Fig. 4E). We speculate that these densities might be chromatin-associating protein complexes such as transcription factors, chromatin remodelers, or their subunits or subnucleosomal particles. Taken together, our *in vitro* DNA-length measurements are consistent with nucleosome-mapping studies.

### Natural chromatin has few ordered positions

The crystal structure of a tetranucleosome and the cryo-EM structures of 30-nm fibers both have zigzag motifs with face-to-face nucleosome packing (Schalch et al., 2005; Song et al., 2014). In yeast chromatin, we found that only a few nucleosomes were packed face-to-face (Fig. 5A) or in a zigzag motif (Fig. 5B). To detect more regular packing motifs, we performed 2-D classification on subtomograms that contained multiple nucleosomes (Fig. 5C). In agreement with our visual inspection, only one class showed face-to-face packing and furthermore, there was no dominant motif (Fig. 5C).

**Figure 5.**
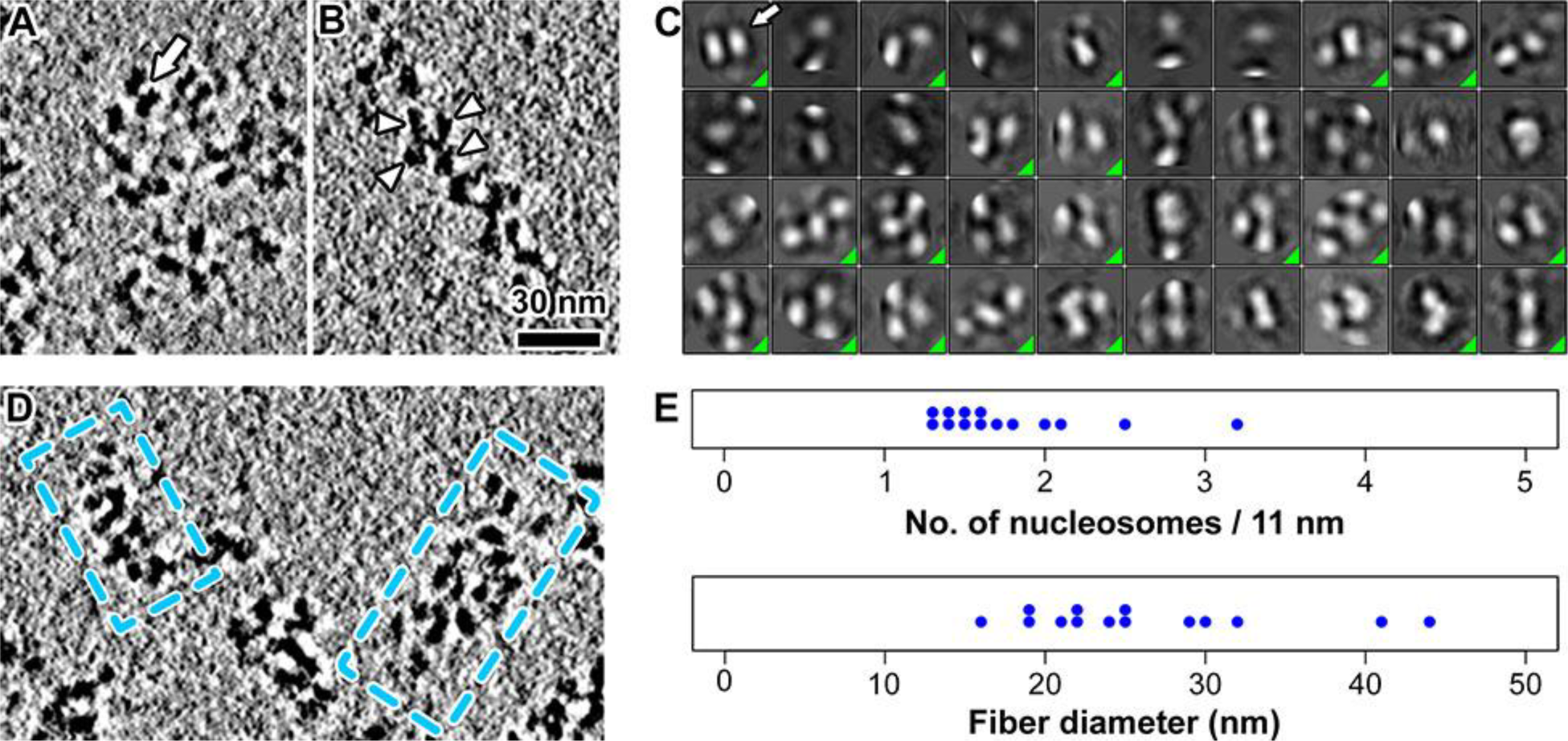
Natural chromatin adopts few regular packing motif. (A) Tomographic slice (10 nm) of yeast chromatin showing face-to-face packing (arrow). (B) Tomographic slice (6 nm) of yeast chromatin showing zigzag motif. Arrowheads: nucleosomes. (C) Two-dimensional classification of yeast lysate in 50 mM EDTA, ordered left-to-right, top-to-bottom from the most abundant to least abundant. Classes that represent actual nucleosome-nucleosome interactions have a green triangle in the lower-right corner. The contrast is inverted relative to the raw tomograms. Arrow: a class showing face-to-face packing. (D) Tomographic slice (10 nm) of yeast chromatin showing irregular 30-nm fibers. (E) Measurement of fiber compaction parameters.

Artificial nucleosome arrays with a 167-bp repeat (identical to the yeast nucleosome repeat length) are organized as compact 30-nm fibers (Routh et al., 2008). To test if natural yeast chromatin fibers (Fig. 5D) are as compact as these artificial 30- nm fibers, we determined the number of nucleosomes per 11 nm in the best-resolved fibers (Fig. 5E). Compared to the artificial fibers, most natural fibers are less compact (∼1.5 vs 6.1 nucleosomes per 11 nm) and have a broader distribution of diameters (15 to 45 nm vs. 15 to 25nm, Fig. 5E) (Routh et al., 2008). The two best-resolved picoplankton 30-nm fibers (Fig. S2G and H) were also loosely packed (3.3 and 2.4 nucleosomes / 11 nm). Therefore, natural chromatin fibers are less compact but much more heterogeneous than artificial fibers.

### Natural chromatin is not very heterogeneous at the nucleosome level

The complexity of natural chromatin could be explained if the nucleosomes are themselves conformationally heterogeneous. To test this hypothesis, we developed a subtomogram analysis workflow that takes advantage of recent advances in 2-D and 3-D image classification (Figs. 6, 7, S9) (Bharat and Scheres, 2016). This approach was able to resolve a few nucleosome conformers that differ in linker DNA orientation. For picoplankton, linker DNA conformation was variable, but the conformers generally fell into three classes (Figs. 6A, S10). In two classes, the linker DNA strands frequently entered and exited the nucleosome core in an “open” instead of “crossed” conformation (Fig. 6A, black and red arrows, respectively; Movie S1). In the third class, only one linker DNA was consistently in the same orientation, and therefore visible. While the 3-D classification in yeast nucleosomes showed higher-resolution features like the DNA gyres of nucleosomes, the linker DNA could not be visualized in most classes except for one class showing a DNA stem-like structure, which might be associated with linker histones (Fig. 7A). Taken together, our 3-D classification reveals that the conformation of nucleosomes is not very heterogeneous, except in the linker DNA.

**Figure 6.**
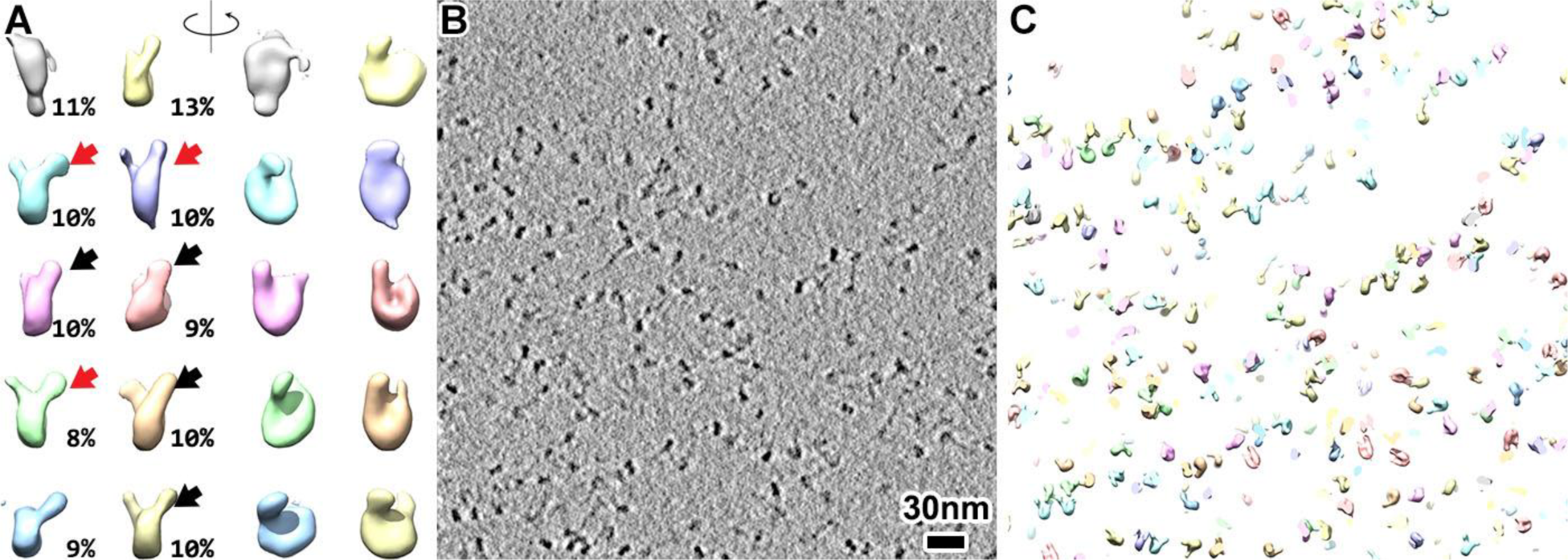
Open zigzag chromatin accommodates open linker DNA. Gallery of nucleosome classes viewed (left) edge on and (right) face on, contoured at 0.5γ to better visualize the linker DNA. The percentage of particles belonging is shown in black numbers at the lower right of each respective class. Black arrows: classes with open linker DNA. Red arrows: classes with crossed linker DNA. Notice that the yellow and blue class only show the density of one linker DNA. (B) Tomographic slice (30 nm) of picoplankton chromatin in lysis buffer with 5 mM EDTA. (C) Three-dimensional classes of template-matched nucleosomes, remapped as a synthetic tomogram.

**Figure 7.**
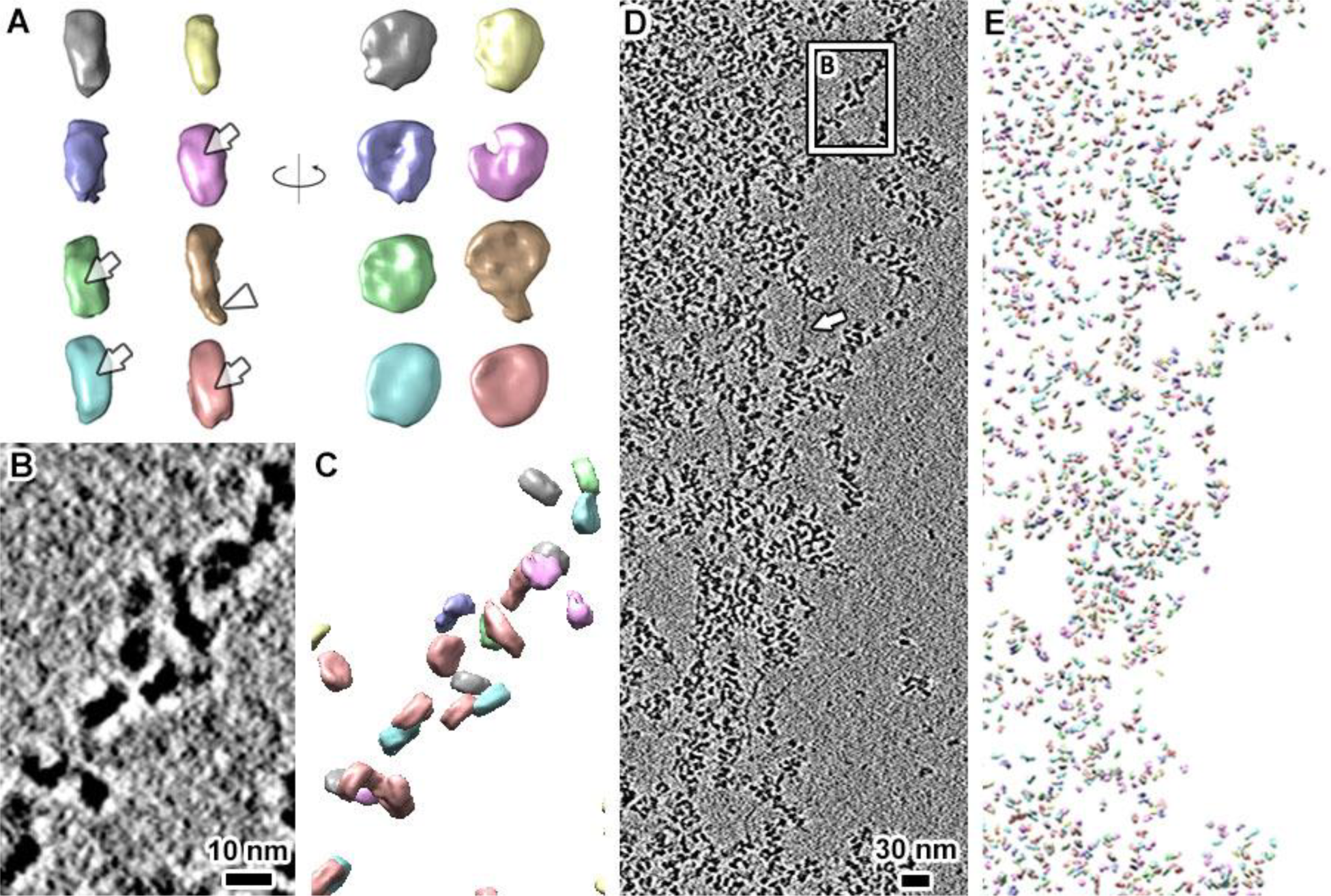
Synthetic tomogram reveals the conformational and configurational heterogeneity of yeast chromatin. (A) Gallery of 3-D classes viewed (left) edge on and (right) face on. Arrows: DNA gyres. Arrowhead: DNA stem-like structure. Note these nucleosome classes are not color coded the same way as in Fig. 4. (B) Five-fold enlargement of an irregular 30-nm fiber boxed in panel D. (C) synthetic tomogram of a 78-nm thick tomogram (same X-Y position as panel C). Arrow: a long stretch of naked DNA. (D) Tomographic slice (10 nm) of yeast nuclei lysed in the presence of 50 mM EDTA. (E) Synthetic tomogram of the same position as panel D).

Even though some of our tomograms reveal the locations of individual nucleosomes, the tomograms are still so crowded and complex that we may have missed some higher-order structures by visual inspection. To further test if the nucleosomes pack into higher-order structures, we remapped the class averages of template-matched nucleosomes into a phantom volume the same size of the original tomogram to make a “synthetic tomogram”. This approach is similar to the work done on polysomes and purified oligonucleosomes (Brandt et al., 2009; Scheffer et al., 2012). Most of the nucleosomes of open zigzags and an example chromatin fiber were visualized in the synthetic tomograms, showing the robustness of our approach (Fig. 6B and C and 7B and C). In both picoplankton and yeast, the nucleosomes do not form long repeating motifs or regular structures of any kind (Fig. 6C and 7E). Furthermore, there were no nucleosome clusters or chains formed by the same conformational class. Therefore, natural chromatin is structurally heterogeneous at the oligonucleosome level.

## DISCUSSION

Our previous cryo-ET studies of picoplankton and yeast challenged the notion that the 30-nm fiber explained most chromatin *in vivo* in unicellular eukaryotes. Here, we have tested how natural chromatin is organized *in vitro*, but using undigested natural chromatin instead of the chromatin fragments found in most *in vitro* studies. This natural chromatin condenses as fibers and masses at much lower divalent-cation concentrations than needed for chromatin fragments (Hansen, 2002; Maeshima et al., 2016). In the absence of divalent cations, picoplankton chromatin decondenses to an open zigzag configuration but yeast chromatin retains an irregular 30-nm fiber configuration. This discrepancy is highlighted in yeast chromatin, which, as discussed below, did not unfold as expected. These observations reveal that natural chromatin is both plastic and unpredictable.

### Picoplankton and yeast chromatin have different sensitivity to divalent cations

Sequestration of divalent cations with EDTA is known to disrupt 30-nm-fiber formation and oligomerization (Widom, 1986). We found that the chromatin of both picoplankton and yeast is more disperse in the absence of divalent cations, but the level of dispersion is different. Picoplankton chromatin adopts an open zigzag configuration whereas the yeast chromatin is much more crowded, with a few irregular 30-nm fibers, even in 50 mM EDTA (Fig. 8). According to an early study (Lowary and Widom, 1989), the yeast chromatin should have become 10-nm filaments. We did not observe such structures, probably due to the sample differences, e.g., purified short yeast chromatin fragments vs. nuclear lysate. Natural picoplankton chromatin therefore follows the divalent-cation-dependent compaction behavior seen in metazoans, whereas yeast chromatin does not.

**Figure 8.**
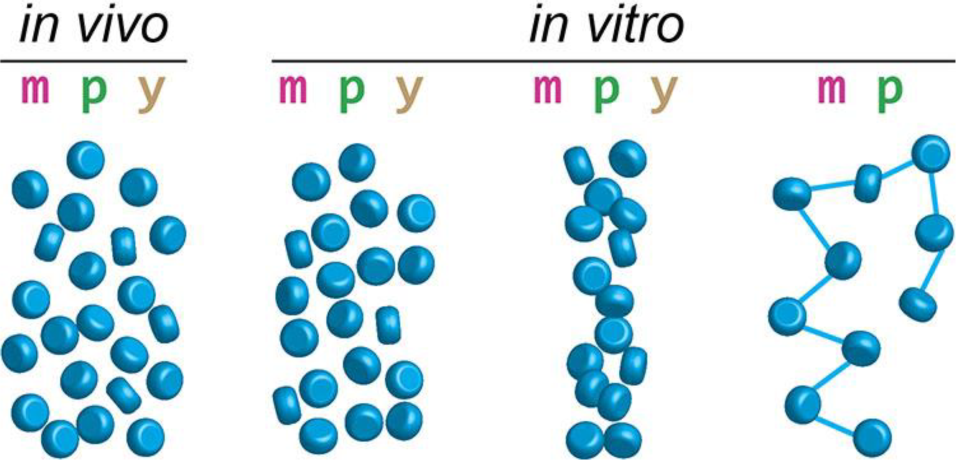
Chromatin structure is highly variable *in vitro*. Nucleosome (blue disc) packing in (m) metazoans, (p) picoplankton, and (y) yeast. *In vivo*, cryo-ET data is most consistent with a chromatin mass. *In vitro,* (left) masses, (middle) irregular 30-nm fibers, and (right) open zigzags have been seen in all three classes of organisms but very rarely in yeast. Linker DNA (blue lines) is depicted only in structures in which it has been observed.

What factors could explain the differences in the higher-order structure of picoplankton and yeast chromatin? First, the nucleosome-repeat length of picoplankton is ∼ 30 bp longer than yeast (Thomas and Furber, 1976; Gan et al., 2013). The shorter yeast linker DNA brings nucleosomes closer together, increasing the chances of interaction between sequential nucleosomes. Second, picoplankton does not have a linker-histone homolog, but yeast does (Downs et al., 2003). Linker histones could keep yeast chromatin compact by, for example, restricting the angles between entering and exiting DNA, which would further disfavor an open zigzag conformation (Thoma and Koller, 1977; Bednar et al., 1998). Third, picoplankton linker DNA is enriched in methylation (Huff and Zilberman, 2014), which stiffens DNA (Nathan and Crothers, 2002). This stiffer linker DNA may keep picoplankton nucleosomes apart as open zigzags in the absence of divalent cations. We hypothesize that linker DNA length and chemistry and the presence of linker histone jointly contribute to chromatin compactability.

### Nucleosomes of natural chromatin pack heterogeneously

Linker DNA has variable length and enters and leaves the nucleosome core particle at variable angles (Yao et al., 1990; Bednar et al., 2017). In the context of undigested chromatin, we also detected linker DNA of differing orientations in the 3-D classes of picoplankton nucleosomes (Fig. 6A). Unlike the cryo-EM study on reconstituted nucleosome arrays (without linker histone) where most nucleosomes had linker DNA in the “closed” conformation (Geiss et al., 2014), we find that picoplankton linker DNA adopts a more “open” conformation, possibly due to the enhanced stiffness of picoplankton linker DNA (see above). In contrast, most 3-D classes of yeast nucleosomes did not show linker DNA, probably due to the crowded packing of nucleosomes that lowers its contrast. We nevertheless saw a stem-like density in one class, similar to a recently published structure of the chromatosome (Bednar et al., 2017).

Beads-on-a-string motifs are characterized by the large ∼ 180° angle made by the linker DNA at the nucleosome entry and exit points (Fig. 1A, structure v). Repetition of this motif maximizes the distance between nucleosomes, resulting in maximum decondensation. These motifs have been routinely seen in dried chromatin that has been adhered to a carbon film (Olins and Olins, 1974; Grigoryev et al., 2009), but very rarely seen in frozen-hydrated chromatin (Bednar et al., 1998; Geiss et al., 2014). We very rarely saw beads-on-a-string in our picoplankton chromatin and did not see a single example in our yeast chromatin; we surmise that if yeast chromatin adapted the beads-on-a-string configuration, we would have observed a structure that could be called a 10-nm filament. These observations are consistent with the crystal structure of the mononucleosome (Luger et al., 1997), from which the linker DNA would emerge at an angle much less than 180° in the absence of large bending forces. Therefore, the most decondensed conformation that natural chromatin can adopt *in vitro* is an open zigzag.

Face-to-face nucleosome packing has been observed in oligonucleosome arrays (Robinson et al., 2006) and isolated starfish sperm chromatin (Scheffer et al., 2012). Here we found that the face-to-face interactions exist in both picoplankton and yeast, but are rare. The face-to-face packing probably involves the well-known interaction between a positively charged histone tail on one nucleosome with the acidic patch on a neighbor (Luger et al., 1997a; Funke et al., 2016). The rarity of face-to-face packing in natural chromatin is at odds with the cryo-EM models of recombinant 30-nm fibers, in which every single nucleosome is involved in a face-to-face packing interaction (Song et al., 2014). In that study, the absence of histone marks, the glutaraldehyde fixation, and the periodic spacing between adjacent nucleosomes could have stabilized the face-to-face packing interaction, thereby explaining why the structures of reconstituted nucleosome arrays differ so much from natural chromatin.

An earlier study found that short nucleosome arrays with a yeast-like nucleosome-repeat length (167 bp) could form ordered, dispersed, 30-nm fibers *in vitro* (Routh et al., 2008). We did not observe such fibers. Instead, most yeast nucleosomes pack into a large mass without any regular motifs; in the few other cases, chromatin packs as irregular 30-nm fibers. The formation of chromatin masses can be explained by the interdigitation of fibers, i.e., inter-fiber interactions force the irregular 30-nm fibers to be in an unfolded state, mixed with the nucleosomes of other unfolded 30-nm fibers (Eltsov et al., 2008). These masses could also be explained by hierarchical loops, which are compatible with more heterogeneous packing than helical models (Grigoryev et al., 2016). More importantly, 30-nm fibers are found at the less-crowded periphery of chromatin mass *in vitro*. This phenomenon supports the idea that the chromatin crowding can prevent fiber formation. However, due to technical limitations, we cannot exclude the possibility that the 30-nm fibers do not unfold, but are instead indistinguishable when closely packed (Grigoryev and Woodcock, 2012).

### Chromatin is more compact *in vitro* than *in vivo*

Our cryo-ET studies of two organisms gives us a rare opportunity to assess the factors that contribute differences in 3-D chromatin organization *in vivo* and *in vitro*. One major difference is that nucleosomes pack slightly closer *in vitro* than *in vivo*, especially in yeast (Fig. 4 H and I). Although our lysis approach is gentle, it cannot keep chromatin remodelers and transcription factors from dissociating from the chromatin. The presence of such factors *in vivo* might prevent the chromatin from folding into a compact higher-order structure. This notion was supported by a study showing that when the Widom 601 nucleosome-positioning element was inserted into yeast genome, both the positioning and the occupancy changed substantially compared to *in vitro* chromatin arrays (Perales et al., 2011). We hypothesize that chromatin remodelers, transcription factors, and transcription itself keep chromatin dispersed *in vivo*; the absence of these factors *in vitro* allows chromatin segments to pack closer together.

## MATERIALS AND METHODS

### Picoplankton cell culture

*Ostreococcus tauri* cells (strain OTH95, Roscoff Culture Collection strain RCC 745) were grown in artificial seawater containing Sigma sea salt and Keller enrichment medium (Table S1) in a 12h:12h light:dark cycle. Lighting was provided by T5050 white light-emitting diodes, passed through a ‘Moonlight blue’ filter (Lee Filters, #183; Panavision, Los Angeles, CA). This setup produced an illuminance of ∼ 400 - 500 lux when measured at the approximate position of the cells, with an Amprobe LM-100 digital light meter (Danaher, Inc., Everett, WA). Cells (30 - 50 ml) grown this way were loosely synchronized and were mostly in mid-G1 phase at the beginning of the light phase (Corellou et al., 2005). The cells were harvested in mid-log phase (OD_600_ ∼ 0.05 - 0.1) shortly after the light-to-dark transition and then pelleted by centrifugation at 5,000 × *g* for 10 minutes at 4°C.

### Picoplankton lysis

Cells were then resuspended in 1 ml of prechilled (4°C) fresh artificial seawater and re-centrifuged at 5000 × *g* for 1 minute. The cell pellet was then resuspended in prechilled lysis buffer, yielding a final OD_600_ ∼ 20. The cells were incubated on ice for either 7 - 9 minutes (1 mM Mg^2+^) or 10 - 15 minutes (0 Mg^2+^ or 5 mM EDTA), measured from when the cells were resuspended in lysis buffer to the point of plunge freezing. Using a combination of sucrose and glycerol, the concentration of soluble particles (ions and undissociated molecules) in the lysis buffer was adjusted to ∼820 mM, which is approximately 4/5 of the concentration of soluble buffer molecules in artificial seawater (∼ 1 M); for details, see Table S2. The picoplankton burst in all lysis buffers. While the plasma membrane was completely ruptured, the contents of the largest two organelles the nucleus and the chloroplast – remained physically associated. This association was fortuitous because the chloroplast remnants, which were high contrast and could be located even at low magnification, facilitated the search for the adjacent chromatin.

### Yeast nucleus isolation and lysis

Nuclei were isolated using the Yeast nuclei isolation kit (Abcam 206997), with modifications. Wild-type yeast (YEF473A; a gift from Kerry Bloom) cells were grown in 30 ml yeast peptone dextrose (YPD) medium to mid-log phase (OD_600_ ∼ 1) in a shaker (30°C, 250 RPM). The cells were pelleted by centrifuging at 3,000 × *g* for 5 minutes at room temperature in 50 ml conical tubes, washed twice by resuspension in 1 ml H_2_O, then re-centrifuged at 3000 × *g* for 1 min. The cell pellet was then resuspended in 1 ml Buffer A (containing 10 mM DTT, from the kit), and then incubated for 10 minutes in a 30°C water bath. Cells were then centrifuged at 1500 × *g* for 5 minutes and then resuspended in 1 ml Buffer B (from the kit) containing lysis enzyme cocktail (diluted 1:1,000). The mixture was incubated in a 30°C shaker for 15 min, and then centrifuged at 1,500 × *g* for 5 minutes at 4°C. The pellet was resuspended in 1 ml prechilled Buffer C (from the kit) with protease inhibitor cocktail. The suspension was transferred to a glass Dounce homogenizer on ice and then the cells were lysed with 15 strokes. The suspension was incubated with shaking for 30 minutes at room temperature and was then centrifuged at 500 × *g* for 5 minutes at 4°C to remove the debris. A small aliquot of the supernatant was stained with 4′,6-Diamidino-2-phenylindole dihydrochloride (DAPI) to verify that the isolated nuclei were abundant. The supernatant was then re-centrifuged at 20,000 × *g* for 10 minutes at 4°C. The nuclear pellet was resuspended in pre-chilled lysis buffer (see Table S2) and incubated for 15 minutes on ice prior to plunge freezing.

### TSA treatment

Wild-type yeast were grown in 30 ml YPD to mid-log phase (OD_600_ ∼ 1) in a shaker (30°C, 250 RPM). Cells were pelleted at 3,000 × *g* for 5 minutes and then resuspended in 30 ml YPD with 16.4 µM TSA (5 mM stock in dimethyl sulfoxide, Sigma T1952). The cells were then incubated for 1 hour at 30°C before nuclei isolation. During nuclei isolation, the solutions used up to the spheroplasting step did not contain TSA. After spheroplasting, all solutions contained 82 µM TSA.

### Plunge-freezing

Colloidal gold (used for fiducial-based tomographic image alignment; see below) has a tendency to aggregate in the presence of seawater unless first treated with bovine serum albumin (BSA) (Gan et al., 2011). Therefore, the colloidal gold (20 nm, BBI solutions, Cardiff, UK) was first suspended in 10 mg/ml BSA, pelleted at 18,000 × *g* for 5 minutes, and then the supernatant was discarded. The gold was treated with BSA twice in this way. The BSA-treated gold was then added to the cells just before plunge-freezing.

Grids (CF-4/2-2C-T and CF-2/2-2C-T, Protochips, Morrisville, NC) were plasma cleaned 90 seconds at 15 mA with a Emitech K100X glow-discharge unit. Treated cells (3 µl) were added to each side of a plasma-cleaned grid. The grids were blotted with filter paper (Whatman, Grade 1) for 1 second, blot force 1, followed by a 5-second wait time, then plunged into 67/33 (% v/v) liquid propane/ethane mixture (Tivol et al., 2008) using a Vitrobot, Mark IV (Thermo, Waltham, MA). The relative humidity in the sample chamber of the Vitrobot was kept at 100%. The grids were then stored in liquid nitrogen.

### Cryo-ET and image processing

Data-collection parameters are detailed in Table S3. Tilt-series alignment and tomographic reconstruction were done using Etomo (Mastronarde, 1997). Because only the positions corresponding to the nuclear contents were of interest and because some sample positions warped during data collection, most tilt-series were aligned using only the gold fiducials in the vicinity of the nuclear densities. Some of the beads were automatically removed with the Etomo “bead eraser” tool. For yeast samples, all tilt series were aligned using patch tracking because of insufficient gold fiducial numbers. For yeast and picoplankton samples treated with 0 mM Mg^2+^ or with EDTA, the tilt series were first corrected for the effects of the contrast-transfer function and then 2-D low-pass filtered with cutoff in the range of 0.2 - 0.35 pixel^-1^ and sigma value of 0.05 pixel^-1^; these settings gently attenuated spatial-frequency components starting at 4.5-nm (picoplankton) to 2.6-nm (yeast) resolution. For picoplankton samples treated with 1 mM Mg^2+^, the contrast was much lower due to sample thickness and crowding effects, necessitating the two-fold binning of the tilt series to improve the visualization. Other parameters were kept as Etomo defaults.

### Naked-DNA length analysis

To facilitate the measurement of naked DNA length, the tomograms were binned by 2 and then low-pass filtered (cutoff=0.3, sigma=0.05). DNA in a tomogram was tracked semi-automatically using the Livewire drawing tool in 3dmod, saved as a model file, then extracted with the IMOD program imodinfo.

### Template matching

Template matching was done using PEET (Heumann, 2016). Two types of templates were used for picoplankton: a featureless cylinder (6-nm thick, 10-nm diameter), generated using beditimg (Heymann and Belnap, 2007); for yeast: a subtomogram of a high contrast nucleosome-like particle was selected. The choice of reference did not influence the subsequent analysis of most particles because a low cross-correlation (CC) cutoff was used (see below) and the rotational alignment solution was discarded prior to further analysis. For *O. tauri*, however, we found that hits corresponding to nucleosomes with the superhelical axis nearly perpendicular to the ice surface were off-center. These nucleosomes were better centered when we used a low-pass-filtered subtomogram average of a RELION 3-D class as a reference (see below). To minimize the effects of neighboring densities, subvolumes were isolated with a thick cylindrical mask with a soft edge. To minimize the number of false negatives, we kept the template-matching hits that were spaced as close as 6 nm, corresponding to face-to-face packed nucleosomes. This minimum distance resulted in many “overlapped” candidate nucleosomes (hits), of which one was removed automatically at each position at the end of the search. To further minimize the number of false negatives, all hits with a CC (relative to the template) greater than 0.2 were saved and then visualized in the original tomogram; the CC values were displayed using the 3dmod “fine grain” feature. The CC cutoff was then manually increased in 0.05 increments until most spurious hits (carbon support edge, ice contaminants, and aggregates) were eliminated. The final filtered hit list was then subject to heterogeneity analysis using RELION (see below).

### Nearest-neighbor distance analysis

For a fair comparison between different datasets, we used the average of CC coefficients of all template-matching hits as cutoff. For example, if the average CC value in dataset A was 0.4 and the one in dataset B was 0.3, then we used 0.4 as cutoff for A and 0.3 for B. We then made small adjustments to the CC cutoff to ensure the low occurrence of obvious false positives (like ribosomes and ice) and false negatives. The coordinates of the nucleosome hits were then imported into Matlab. NND and 10^th^ NND were calculated using the Matlab function nearestneighbour.m. The script is available upon request.

### Heterogeneity analysis and subtomogram averaging

Subtomograms were analyzed using the subtomogram-averaging and classification routines (Bharat et al., 2015; Bharat and Scheres, 2016) as implemented in RELION 1.4 and 2.0 (Kimanius et al., 2016; Scheres, 2012a, b). Because CTF phase flipping was already done in IMOD, we disabled phase flipping in RELION. We also disabled the dose-weightage scheme because the images in the second half of the tilt series were severely underweighted due to the large cumulative dose. The missing-wedge effects were compensated for by the RELION CTF model. Candidate nucleosome positions were imported into RELION. Owing to the crowdedness of yeast nucleosomes, we used a smaller box (24 nm) and a smaller mask (17-nm diameter) for classification to exclude most densities of adjacent nucleosomes. Each nucleosome was averaged along the tomographic Z axis to produce a pseudo projection. These projections were then subjected to 2-D classification (50 or 100 classes). “Junk particles” such as ice, contaminants, or carbon-support features were manually removed. If the identity of the class ("junk” vs. real nucleosome) was ambiguous, the member particle positions were visualized in the context of the original tomogram, allowing for discrimination of junk and real particles. Our decisions to exclude/include ambiguous classes were guided by biologically meaningful features such as the size and shape of the density and the proximity to the other nucleosomes (chromatin-associated nucleosomes are not found “floating” alone). The remaining good particles were subjected additional rounds of 2-D classification and elimination of junk classes until only reasonable nucleosome-like classes remained.

To better discriminate between nucleosome conformational states, we performed 3-D classification without the use of an external reference. New junk classes were found and removed at the end of each round of classification. We found that the number of classes decreased as we increased the resolution used for classification to the Nyquist limit. This resolution dependency is probably due to the lower signal-to-noise ratio of the higher-resolution data. Starting with 20 classes and using data to the Nyquist limit (20 Å for yeast), fewer than 10 distinctive conformational classes remained at the end of 2 - 3 sequential classification-and-junk-removal rounds. These classes were then subjected to synthetic tomogram construction. For picoplankton chromatin in 1 mM and 0 mM Mg^2+^, we also attempted 3-D classification of higher-order structures by increasing the mask size, but found that most of the classes did not converge to anything biologically meaningful, probably due to the extreme heterogeneity at the level of oligo-nucleosome structures. Our 2-D classification of picoplankton chromatin in 5 mM EDTA failed to find any convincing face-to-face or other abundant higher-order motifs because most of the nucleosomes were too far apart to interact.

### Synthetic tomogram construction

Template-matching hits that were subjected to the 2-D and 3-D classification process were mapped back into a phantom volume the size of the original tomogram using the EMAN2 program e2proc3d (Tang et al., 2007). The script to do this is available upon request.

### Figures

Density maps were rendered with UCSF Chimera (Pettersen et al., 2004).

### Data sharing

Tilt series data of all samples presented in this paper have been made accessible in the EMPIAR online database: (EMPIAR-10098). One example tomogram each of picoplankton and yeast has been deposited in the EMDB (EMD-6737).

## Acknowledgements

We thank the CBIS microscopy staff for support and training. We thank François-Yves Bouget for advice of Ostreococcus cell-culture conditions, Rado Danev for suggesting 2-D classification of subtomograms, and Kerry Bloom for yeast strains. We also thank the Gan team and David Shore for discussions. SC, YS, CC, and LG were supported by NUS startups R-154-000-515-133, R-154-000-524-651, and D-E12-303-154-217, a MOE T2 R-154-000-624-112, and a NUS YIA R-154-000-558-133.

## Contributions

SC - experiments, project design, writing; YS - experiments; CC - experiments, editing; JS - training; LG - experiments, project design, writing.

